# A Single-Cell Atlas of the Breast Cancer Microenvironment Identifies Subtype-Specific Tumor-Immune Landscapes and Vulnerabilities

**DOI:** 10.1101/2025.09.26.678647

**Authors:** Liron Zisman Schachter, Asaf Pinhasi, Keren Yizhak

## Abstract

Breast cancer is a heterogeneous disease, with prognosis and treatment shaped by subtype and stage. In this study we integrated 31 single-cell RNA-seq datasets, totaling 1.2 million cells from 376 samples. Our analysis revealed distinct subtype- and stage-specific TME states, and in both TNBC and Luminal A, we uncovered connections between tumor-intrinsic programs and the microenvironment, directly linked to patient survival and therapeutic response. In TNBC, tumor programs differed by stage, with primaries linked to immune-modulatory states and metastases to poor survival. These programs showed elevated TROP2 expression, suggesting sensitivity to TROP2-targeted therapies. Stage-specific checkpoints linked tumor programs to PD-1/LAG3 in primaries and CTLA-4 in metastases. In Luminal A, ER⁺ tumors showed proliferative signaling linked to favorable endocrine therapy response but also immune exclusion, whereas ER⁻ tumors exhibited MYC-driven programs with poor prognosis, yet higher immune infiltration. Our breast cancer atlas outlines evolving therapeutic vulnerabilities, providing a framework for precision medicine and clinical trials design.

## Introduction

Breast cancer continues to be the most frequently diagnosed cancer among women worldwide and remains a leading cause of cancer-related mortality^1^. In the United States alone, it accounts for approximately 30% of all new cancer diagnoses among women, with an estimated 316,950 new cases and 42,170 deaths in 2025^2^. Notably, early detection through supplemental MRI or ultrasound screening, particularly for high-risk individuals such as those carrying BRCA mutations, has significantly improved outcomes by identifying the disease at earlier and more treatable stages^3,2^.

Breast cancer is a highly heterogeneous disease, encompassing distinct molecular subtypes that guide prognosis and therapeutic decision-making^4^. The primary subtypes, Luminal A (LA), Luminal B (LB), HER2-enriched, and TNBC, differ significantly in terms of biology, treatment response, and clinical outcomes^6^. Each subtype presents unique challenges in both localized and metastatic settings, driving the need for tailored therapeutic approaches^4^.

LA is characterized by high expression of estrogen and progesterone hormone receptors and low proliferation indices. This group often responds well to endocrine therapy, such as tamoxifen or aromatase inhibitors. Recent FDA-approved therapies include selective estrogen receptor degraders (SERDs) and cyclin-dependent kinase (CDK4/6) inhibitors that further enhance endocrine therapy efficacy. In advanced disease, CDK4/6 inhibitors have demonstrated an overall survival (OS) benefit of approximately 12–15 months compared to endocrine therapy alone^5,6^. The duration of response to endocrine therapy in LA patients can extend beyond two years in many cases, particularly when combined with CDK4/6 inhibitors. However, most patients eventually develop resistance and cease to respond, highlighting the need for predictive biomarkers and alternative treatment strategies^7^. Recurrence rates for LA after local excision range from 10–15%. However, with appropriately tailored adjuvant therapies adjusted based on tumor characteristics, the risk of recurrence can be reduced to as low as 1–5%^8–10^.

Luminal B breast cancer shows higher recurrence and more aggressive behavior than Luminal A. Although treatment typically involves both endocrine therapy and chemotherapy, the risk of recurrence remains a significant clinical challenge^11,12^. HER2-enriched tumors have high recurrence rates (30–50%) after local excision but have seen improved outcomes with HER2-targeted therapies. However, despite advances like antibody-drug conjugates, optimizing long-term control and reducing relapse rates in both subtypes continue to pose major therapeutic hurdles^13,14^.

Unlike LA, LB, and HER2-enriched subtypes, TNBC remains the most difficult to treat, showing the least progress in improving overall survival and achieving durable responses. Recurrence rates after local excision are high (30–40%), and even with adjuvant chemotherapy, they remain around 20%^15,16^. In metastatic disease, despite recent advances with immune checkpoint inhibitors and antibody-drug conjugates, overall survival improvements are modest, typically 4–6 months, highlighting the urgent need for more effective, durable treatment strategies^17,18^.

Recent advances in scRNA-seq have provided unprecedented insights into the tumor microenvironment (TME), revealing the heterogeneity of cellular populations within and across breast cancer subtypes. These findings challenge traditional subtype classifications and underscore the complexity of the TME^19^. Notably, scRNA-seq has shed light on the classification of tumors as “hot” (immune-inflamed) or “cold” (immune-deserted or excluded), a framework increasingly used to predict response to immunotherapy^20^.

To decipher how molecular and cellular heterogeneity within the breast TME shapes clinical behavior, we performed the most comprehensive to date, integrative analysis of scRNA-seq data. Focusing on subtype-specific TME features and therapeutic vulnerabilities, our results reveal a dynamic shift between “cold” and “hot” microenvironments across disease states. In localized disease, TNBC tumors were highly infiltrated by exhausted T lymphocytes, whereas Luminal A tumors were largely, dominated by anti-inflammatory macrophages and secretory fibroblasts that fostered immune exclusion. In metastatic settings, LA tumors evolved towards a more inflamed state, marked by a shift towards pro-inflammatory macrophages and a reduction in estrogen and progesterone receptor expression. These changes were associated with MYC-high transcriptional states, endocrine therapy resistance, and poor clinical outcomes. Moreover, in TNBC, we observed stage-dependent checkpoint profiles, with PD-1 and LAG-3 predominating in primary tumors and CTLA-4 in metastases. Additionally, EMT-associated programs emerged in metastatic TNBC, characterized by chromosomal amplifications and increased TROP2 expression, suggesting a potential vulnerability for targeted therapy. Together, these findings illuminate how subtype- and stage-specific cellular ecosystems evolve across disease progression, providing a foundation for precision therapeutic strategies and immunotherapy trials. Our curated BC atlas has been made public through an online repository at: https://breastscape.twix.technion.ac.il

## Results

### Cohort Composition and Dataset Integration

To construct a comprehensive single-cell atlas of the breast TME, we aggregated scRNA-seq from 376 samples derived from 263 patients across 31 publicly available datasets (Figure 1A, Supplementary Table 1, Methods) ^20–49^. The cohort included 239 samples from primary breast tumors, 37 originated from adjusted affected lymph nodes and 23 from distant metastatic lesions.

**Figure 1.**
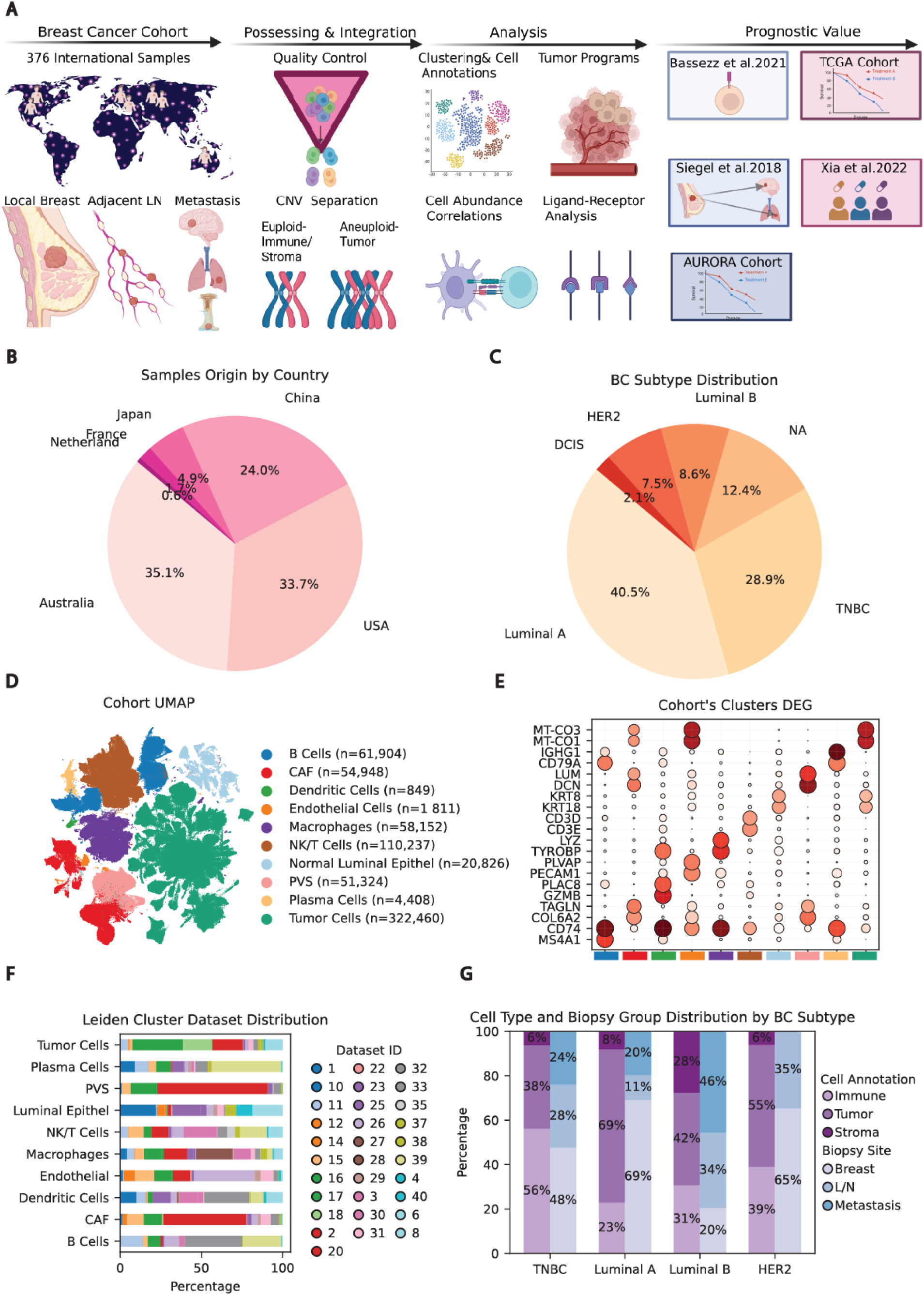
Overview of study design, cohort characteristics, and data structure. **(A)** A schematic representation of the study workflow. **(B)** Geographic distribution of cohort samples by country. **(C)** Distribution of breast cancer subtypes across the cohort. **(D)** UMAP visualization of the combined cohort, colored by major cell types. **(E)** Dot plot showing the expression of top DEGs for each cluster. **(F)** Cohort’s clusters dataset distribution **(G)** Breakdown of sample collection sites and corresponding cell types by breast cancer subtype.

The rest originated from normal breast tissue, DCIS or unknown location samples. In total, 686,919 cells, all sequenced using 10x droplet-based scRNA-seq platforms, passed our QC process. Each dataset was processed independently through a standardized pipeline to ensure high data quality and consistency (Methods). The aggregated datasets reflect substantial geographical diversity, underscoring their broad applicability, with studies conducted in Australia (35.1%), the United States (33.7%), China (24.0%), Japan (4.9%), France (1.7%), and the Netherlands (0.6%, Figure 1B). Among the annotated samples, LA was the most prevalent subtype, representing 40.5% of the integrated cohort, followed by TNBC at 28.9%, and LB at 8.6%. HER2-enriched and ductal carcinoma in situ (DCIS) subtypes accounted for 7.5% and 2.1% of the cohort, respectively, while 12.4% of samples lacked subtype annotation (Figure 1C).

To distinguish aneuploid tumor cells from euploid stromal and immune cells, we performed copy number variation (CNV) analysis (Methods) ^50^. Then, based on clustering and differentially expressed genes (DEG) ^51^, stromal and immune cells were identified, enabling the classification of cells into tumor, immune, and stromal compartments (Methods). This integrative approach resulted in three high-resolution cellular groups comprising 322,460 tumor cells, 235,550 immune cells, and 128,909 stromal and normal epithelial cells, providing a comprehensive resource for downstream analysis of the breast cancer TME (Figure 1D-E; Methods; Supplementary Table 2). To ensure balanced representation across datasets, we quantified the proportion of cells each cohort contributed to each cluster. This analysis confirmed an even representation, validating the robustness of our integration and clustering approach (Figure 1F-G).

### Divergent Immune Infiltration in Primary and Metastatic Breast Cancer

It is well established that TNBC exhibits the highest levels of immune infiltration, whereas Luminal A tumors display the lowest, an observation that was consistently recapitulated in our dataset (Figure 2A)^52^. However, our dataset allows comparison not only across breast cancer subtypes but also across disease stages. Investigating the abundance of immune infiltration across different breast cancer subtypes revealed distinct patterns between localized and metastatic disease. In localized breast cancer, TNBC exhibited the highest levels of immune infiltration, followed by HER2-enriched, Luminal B, and Luminal A subtypes (all pairwise comparisons, p < 0.05; two-sided Wilcoxon rank-sum test; Figure 2B). However, in metastatic disease, no significant differences in immune infiltration were observed among TNBC, Luminal A, and Luminal B subtypes, although substantial inter-sample variability was noted (Figure 2C).

**Figure 2.**
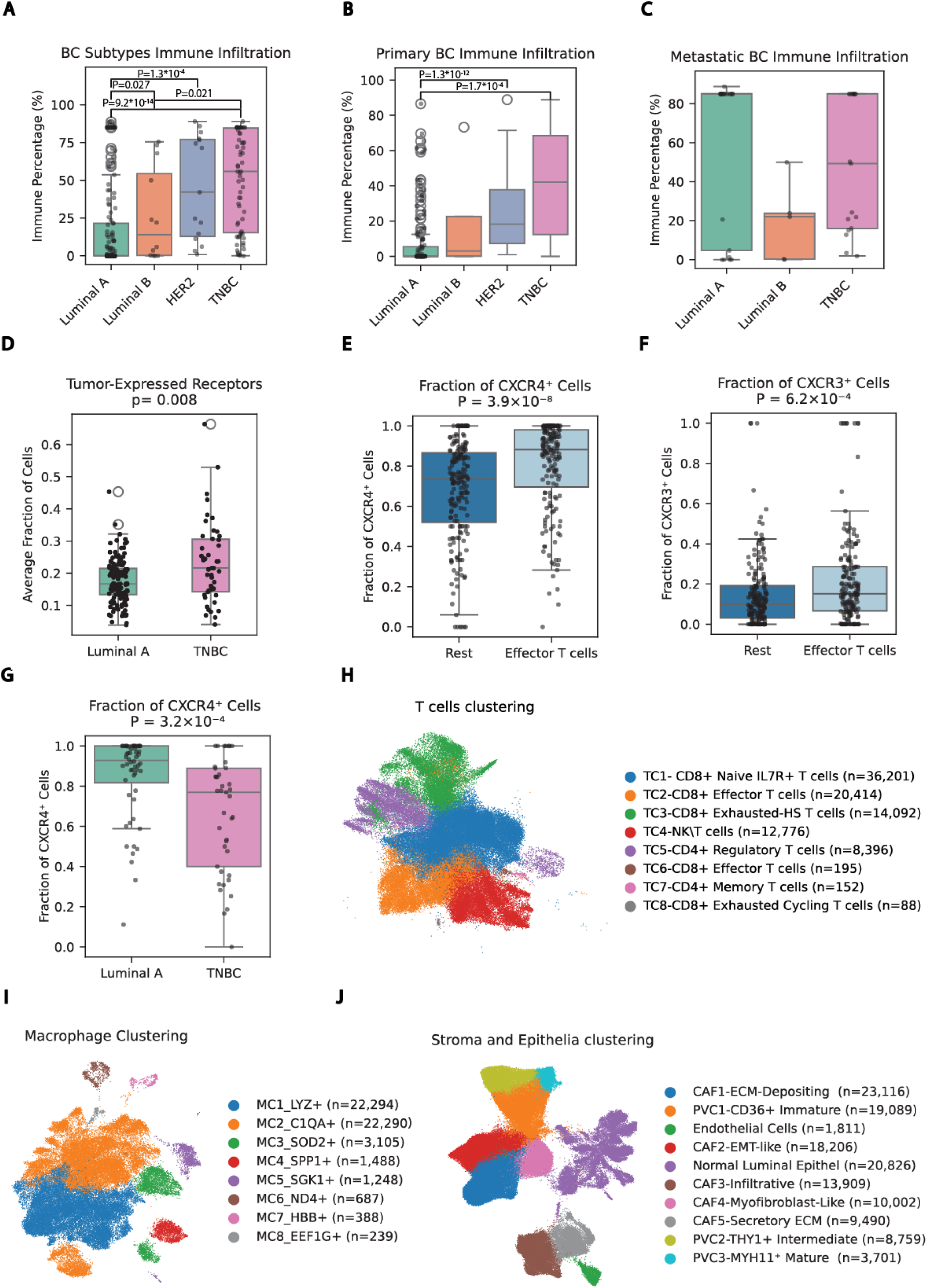
Immune infiltration, cell–cell communication, and TME sub-clustering across breast cancer subtypes. **(A–C)** Percentage of immune cell infiltration in breast cancer subtypes across (A) all samples (B) primary tumors and (C) distant metastatic sites. **(D)** Mean fraction of tumor cells expressing: CXCR4, ITGB1, SDC1, SDC4, VASP in LA vs. TNBC samples **(E-F)** Fraction of (E) CXCR4+ (F) CXCR3+ cells in Effector T cells vs. all other T cells clusters **(G)** Fraction of CXCR4+ cells in effector T cells of LA vs. TNBC samples **(H)** UMAP of T cells, colored by clusters annotations. **(I)** UMAP of macrophage populations, colored by clusters annotations. **(J)** UMAP of stromal cells, colored by clusters annotations.

To further explore the cold-to-hot TME paradigm, we conducted a ligand–receptor (L–R) analysis focused on chemokine signaling interactions among tumor, immune, and stromal cells (Methods). Among the significant L–R pairs identified, the TNBC TME demonstrated a predominance of tumor-promoting interactions (56.66%), followed by tumor-suppressing (26.66%) and dual/undefined-role interactions (16.68%, Supplementary Figure 1A, Supplementary Table 3). Most interactions involved stromal ligands and tumor-expressed receptors, such as CXCR4, ITGB1, SDC1, SDC4, VASP^53–57^, known to promote tumor progression. To validate these observations, we assessed the average fraction of tumor cells expressing these receptors and found them to be significantly upregulated in TNBC tumor cells compared to LA (p = 0.008, two-sided Wilcoxon rank-sum test; Figure 2D).

A similar analysis in LA samples highlighted a predominance of tumor-suppressing interactions (48.27%), with tumor-promoting and dual/undefined interactions comprising 34.48% and 17.24% of total interactions, respectively (Supplementary Figure 1B; Supplementary Table 3). Notably, this pattern intensified when comparing primary to metastatic LA lesions: primary tumors showed 66.7% tumor-suppressing interactions while metastatic sites shifted to just 14.3%, with tumor-promoting interactions rising to 60% (Supplementary Figure 1C,D; Supplementary Table 4). These observations led us to focus on two chemokine receptors, CXCR4 and CXCR3, as they were frequently involved in the significant interactions between stromal and immune cells. In addition, their expression in both innate and adaptive immune populations has been associated mainly with tumor suppression^58–62^. Among immune cells populations, dendritic cells exhibited the highest expression of CXCR3, followed by T cells, while CXCR4 was predominantly expressed in T, B and dendritic cells (p < 0.05 for all comparisons, two-sided Wilcoxon rank-sum test). Within the T cell compartment, these receptors had the highest expression in the CD8^+^ effector T cell clusters (p = 3.9×10⁻⁸ and p = 3.8×10⁻⁴ for CXCR3 and CXCR4, respectively; two-sided Wilcoxon rank-sum test; Figure 2E,F). In addition, CXCR4 expression was significantly higher in LA samples compared to TNBC (p = 3.2×10⁻⁴; two-sided Wilcoxon rank-sum test; Figure 2G), while CXCR3 expression did not differ significantly across breast cancer subtypes.

Altogether, our findings suggest that TNBC tumors are characterized by a chemokine-driven, tumor-promoting microenvironment dominated by stromal-to-tumor signaling. In contrast, localized LA tumors display a more immune-supportive chemokine landscape, partly mediated by differential expression of key receptors such as CXCR4 and CXCR3. However, this supportive landscape is progressively lost in metastatic LA, where we observed a shift toward tumor-promoting stromal interactions alongside increased immune infiltration.

### Dissecting Immune and Stromal Complexity in the Breast TME

To further resolve the heterogeneity within the breast TME, we clustered the cells in the immune and stromal compartments, separately (Methods)^53^. Overall, we identified 8 macrophage\T cells clusters and 10 distinct stromal clusters. In the T cell compartment, we identified 8 clusters (Figure 2H, Supplementary Table 5, Supplementary Figure 2A,B). TC1 included CD8⁺ naive IL7R⁺ T cells expressing CCR7 and TCF7; TC2 represented Effector T cells expressing NKG7 and GZMK; TC3 contained CD8⁺ Exhausted-heat shock T cells expressing HAVCR2 and HSPA1B; TC4 was composed of CD8⁺ NK/T cells expressing KLRD1 and GZMH; TC5 included CD4⁺ regulatory T cells expressing TIGIT and CTLA4; TC6 captured CD8⁺ effector T cells expressing GNLY and GZMB; TC7 included CD4⁺ memory T cells expressing SELL and LEF1; and TC8 comprised of CD8⁺ exhausted T cells with heat shock and cycling features, expressing HSPB1 and CCDC85B.

Within the macrophage compartment, clusters were annotated based on their top five DEGs, reflecting both TME influences and tissue of origin (Figure 2I, Supplementary Table 6, Supplementary Figure 3A,B). MC1 consisted of LYZ⁺ macrophages expressing FCN1 and LST1; MC2 represented C1QA⁺ macrophages expressing C1QB and C1QC; MC3 included SOD2⁺ macrophages expressing CTSL and FTH1; MC4 captured SPP1⁺ macrophages expressing FTL and CSTB; MC5 was composed of SGK1⁺ macrophages expressing CD74 and C1QC; MC6 consisted of ND4⁺ macrophages expressing ND4L and CCL4; MC7 contained HBB⁺ macrophages expressing PLAC8 and VCAM1; and MC8 comprised APOD⁺ macrophages expressing MUCL1 and NME2.

Clustering of stromal cells revealed 10 distinct populations (Figure 2J, Supplementary Table 7, Supplementary Figure 4A,B): one cluster of normal luminal epithelial cells expressing GATA3; five cancer-associated fibroblast (CAF) clusters; three perivascular cell (PVC) clusters; and one cluster of endothelial cells expressing PECAM1 and VWF. The PVC clusters were defined by maturation states, with high concordance to previously published datasets^63^. PVC1 represented CD36⁺ immature PVCs; PVC2 consisted of intermediate THY1+ cells expressing integrin genes; and PVC3 captured MYH11⁺ mature PVCs with contractile properties. The CAF clusters were annotated based on their DEGs: CAF1 represented ECM-depositing fibroblasts expressing FBLN1 and FBLN2; CAF2 consisted of EMT-like fibroblasts expressing DLK1 and SOX4; CAF3 included infiltrative fibroblasts expressing LGALS3 and MGP; CAF4 contained myofibroblast-like cells expressing MYL9 and TAGLN; and CAF5 captured secretory ECM fibroblasts expressing FAP and COL1A2.

To delineate the diversity in the tumor compartment we applied a soft clustering approach to identify tumor expression programs (TP, Methods). This analysis yielded 30 distinct programs (Supplementary Table 8), each demonstrates the activity level of a defined group of genes across all the cohort’s tumor cell. Overall, this high-resolution characterization of the breast TME enabled us to investigate the interplay between these cellular compartments and their role in disease progression and clinical outcome.

### Stratification of LA by ER Status Reveals Prognostic and Immunologic Differences

As hormone receptor status is a major determinant of treatment strategy in luminal A breast cancer, we next examined to what extent this feature is preserved in metastases. We observed a marked reduction in both estrogen receptor (ER) and progesterone receptor (PR) RNA expression in metastatic LA tumor samples compared to local tumors (P < 1 × 10⁻¹⁶, two-sided Wilcoxon rank-sum test, Supplementary Figure 5A,B). This reduction was evident not only in overall expression levels but also in the proportion of ER and PR positive cells (P < 0.017, P < 0.034 for ER- and PR-positive cell, respectively, two-sided Wilcoxon rank-sum test, Figure 3A, Supplementary Figure 5C).

**Figure 3.**
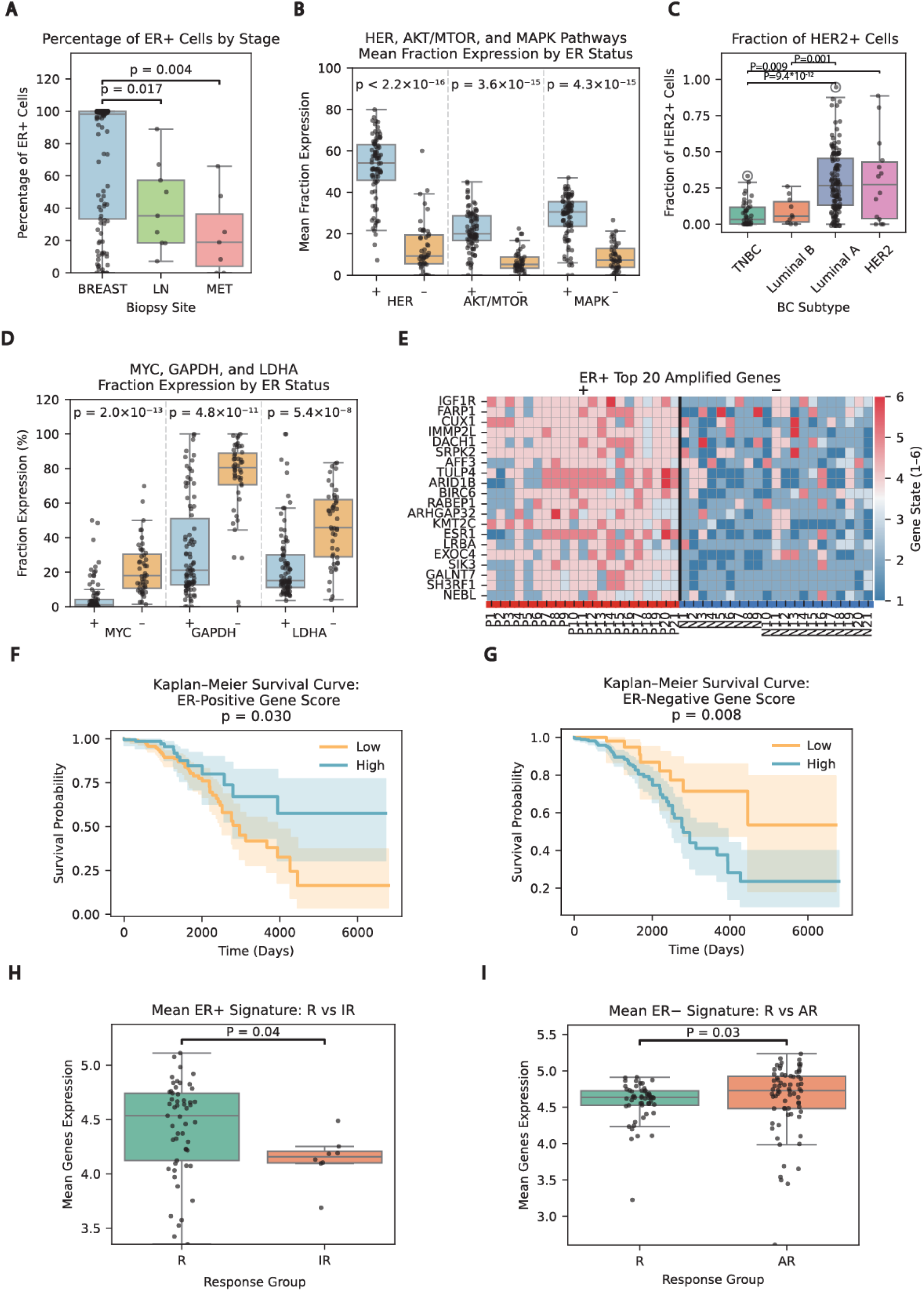
Luminal A ER-dependent populations determine survival patterns and endocrine resistance. **(A)** Percentage of ER-positive tumor cells by biopsy site. **(B)** Mean fraction of cells expressing HER2-4, AKT/MTOR, and KRAS–BRAF–CERK gene signatures in ER-positive (+, blue) versus ER-negative (−, orange) tumor cells. **(C)** Fraction of HER2-positive tumor cells across breast cancer subtypes. **(D)** Fraction of cells expressing of MYC, GAPDH, and LDHA in ER-positive (+) versus ER-negative (−) tumor cells. **(E)** Top 20 amplified genes in ER-positive (P1-21) versus ER-negative (N1-21) CNV clusters. **(F-G)** Kaplan–Meier survival analysis for ER+ BC tumors from the TCGA, stratified by high versus low expression of (F) ER^+^ (high>25%, low<25%) and (G) ER^−^ signatures (high>median, low<median). **(H-I)** Expression of (H) ER^+^ and (I) ER^−^ signatures in responders versus resistant patients to endocrine therapy.

Differential gene expression analysis between ER-positive and ER-negative tumor cells revealed activation of distinct biological programs (Supplementary Table 9). ER-positive tumor cells showed upregulation of well-established proliferation and signaling pathways, including JAK– STAT, KRAS–BRAF–MEK–ERK, HER2/3/4, and mTOR (P < 3.6 × 10^−15^, two-sided Wilcoxon rank-sum test, Figures 3B). Notably, ERBB2 was significantly upregulated in ER-positive cells, despite not being clinically classified as HER2 positive by IHC or FISH. Furthermore, we found no statistical difference between ERBB2 expression in IHC-defined HER2 samples compared to LA samples (Figure 3C). These findings suggest that HER2-targeted therapies may be effective in a subset of LA tumors lacking classical HER2 amplification, a hypothesis supported by recent clinical trials demonstrating benefit from HER2 blockade in this setting^64,65^. In contrast, ER-negative tumor cells appear to be a homogenous population (Supplementary Figure 5D) enriched for metabolic signatures, including anaerobic metabolism (GAPDH, LDHA; P = 4.8 × 10⁻¹¹, P = 5.4 × 10⁻⁸, respectively; two-sided Wilcoxon rank-sum test) and elevated expression of the oncogene MYC (P = 2 × 10⁻¹³, two-sided Wilcoxon rank-sum test, Figure 3D). These pathways are associated with aggressive phenotypes and poor prognosis in breast cancer and other malignancies^66–71^. To our knowledge, this represents the first detailed characterization of ER-negative human tumor cells within LA breast cancer, highlighting their unique metabolic vulnerabilities.

To further investigate these transcriptional differences, we derived ER^+^ and ER^−^ signature by averaging the top DEGs respectively, for each group (Supplementary Table 9, Methods). Analysis of CNV revealed that ∼25% of the genes from the ER^+^ signature, were amplified specifically in ER-positive CNV clusters (Figure 3E, Supplementary Figure 6, Supplementary Table 10, Methods). In contrast, no such amplifications were detected among the genes comprising the ER⁻ signature in ER-negative CNV clusters. Together, these findings suggest that ER-positive cells may sustain their transcriptional programs through genomic amplification, while ER-negative cells exhibit distinct expression patterns including MYC.

To study the applicability of these signatures in bulk setting, we analyzed the BC cohort available in The Cancer Genome Atlas^72^, focusing on ER+ patients. Interestingly, we found that patients with a high ER^+^ signature exhibited significantly better overall survival compared to those with lower signatures (P = 0.03, log-rank test, Figure 3F, Methods). In contrast, a high ER^−^ signature was associated with poor prognosis (P = 0.008, log-rank test, Figure 3G). To ensure that the prognostic value of the ER⁺ signature was not solely attributable to ESR1 expression, we recalculated the signature excluding ESR1. The revised signature remained significantly associated with overall survival (P = 0.05, log-rank test, Supplementary Figure 7). Furthermore, to determine whether these signatures provide prognostic value independent of established clinical covariates, we performed multivariable Cox proportional hazards regression analyses. For both ER^−^ (concordance index of 0.854, SE = 0.028) and ER^+^ signatures (concordance index of 0.851, SE = 0.029), the models remained significantly associated with overall survival after adjusting for age, stage, menopause status, ethnicity, and surgical margin status (Methods, Supplementary Tables 11,12).

Finally, to test the relevance of the ER signature in predicting response to endocrine therapy (ET), we analyzed an independent dataset from Xia et al., 2022^73^ (Methods), which profiled ER-positive breast cancer patients treated with ET. Patients were stratified into responders, acquired resistance, and intrinsic resistance groups. Scoring patients by the ER signature we found that the ER^+^ signature was significantly higher in responders compared to intrinsic resistance patients (P = 0.04, two-sided Wilcoxon rank-sum test, Figure 3H). Conversely, the ER^−^ signature was significantly higher in acquired resistant patients (P = 0.03, two-sided Wilcoxon rank-sum test, Figure 3I) compared to responders. Altogether, these results reveal that ER-positive and ER-negative tumor cell states within LA breast cancer are not only transcriptionally and genomically distinct, but also carry prognostic and therapeutic relevance with potential implications for precision endocrine therapy.

### Linking ER-Driven Tumor States to the Stroma and Immune Microenvironment

To investigate the relationship between LA tumor states and the TME, we assessed ER⁺ and ER⁻ transcriptional signatures in relation to immune infiltration. As shown in Figure 2B,C, immune infiltration patterns differed between primary and metastatic lesions. Here we observed that the ER⁺ signature was significantly enriched in tumors with low immune infiltration, whereas the ER⁻ signature was preferentially associated with tumors exhibiting high immune infiltration (P < 2.2 × 10⁻¹⁶ for both comparisons, two-sided Wilcoxon rank-sum test; Figure 4A).

**Figure 4.**
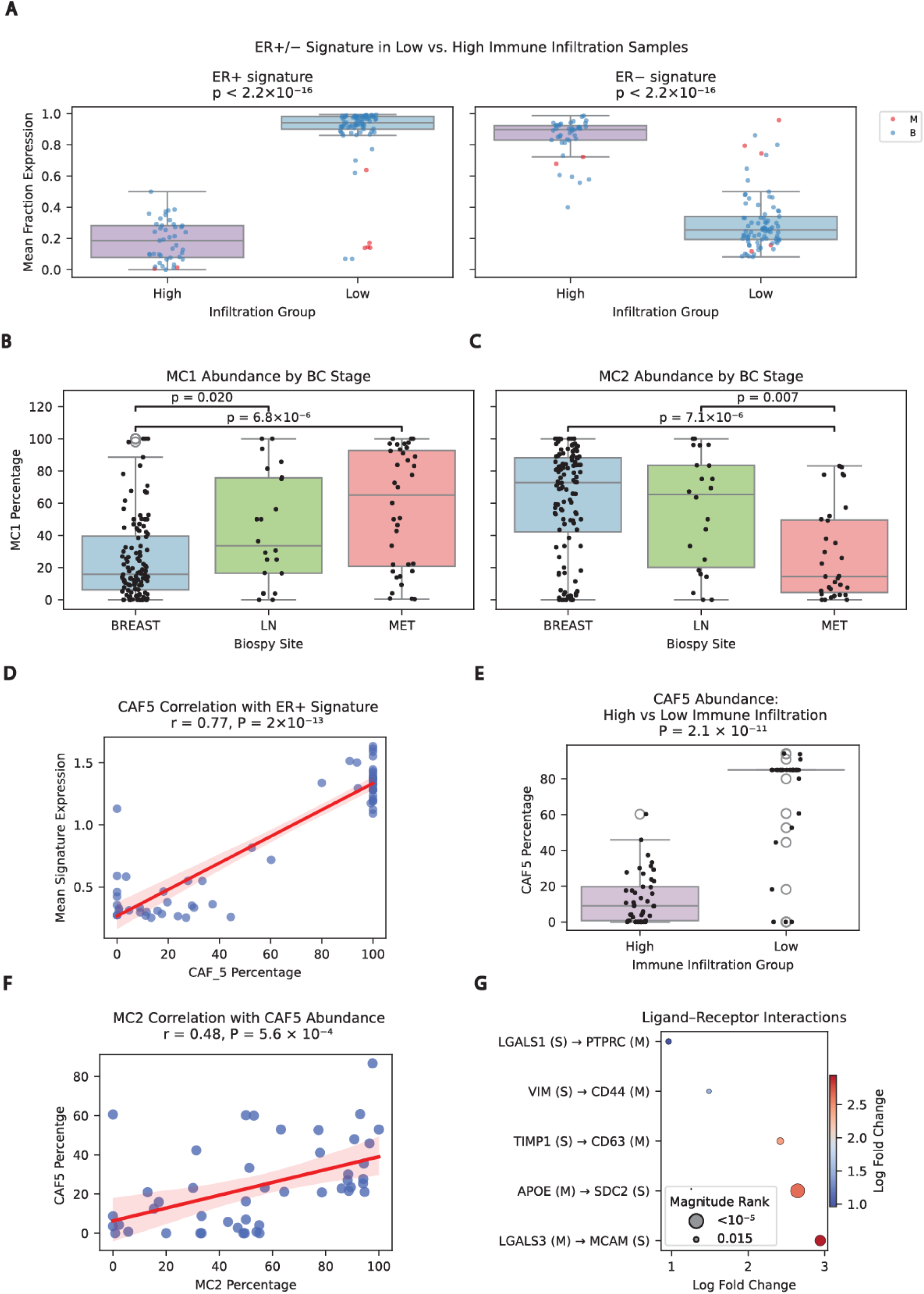
ER-dependent tumor programs shape immune and stromal interactions in Luminal A breast cancer. **(A)** Enrichment of ER positive (left) and ER negative (right) transcriptional signatures in tumors with high versus low immune infiltration. **(B-C)** Abundance of macrophage subclusters MC1 (B) and MC2 (C) across different breast cancer biopsy sites. **(D)** Correlation between CAF5 abundance and ER^+^ signature. **(E)** Abundance of CAF5 in LA tumors with high versus low immune infiltration. **(F)** Correlation between CAF5 and macrophage cluster-MC2 abundance. **(G)** Ligand–receptor interactions between CAF5 and MC2 macrophages.

Further characterization of the immune and stromal compartments revealed striking differences in myeloid cell composition. Cluster abundance analysis demonstrated distinct shifts in macrophage subpopulations between LA breast cancer samples depending on biopsy location (local: breast or adjacent lymph node; metastatic: distant tumor sites). Specifically, MC1–LYZ⁺ macrophages were enriched in metastatic lesions (P = 6.8 × 10⁻⁶, two-sided Wilcoxon rank-sum test; Figure 4B), whereas MC2–C1QA⁺ macrophages were significantly more abundant in primary lesions (P = 7.1 × 10⁻⁶, two-sided Wilcoxon rank-sum test; Figure 4C), suggesting a polarization shift associated with disease progression. MC1–LYZ⁺ macrophages expressed pro-inflammatory and potentially anti-tumor genes, including LYZ, FCN1, TNFSF13B, and IL1B. In contrast, MC2– C1QA⁺ macrophages displayed transcriptional profiles enriched for anti-inflammatory and immunosuppressive markers such as TGFBI, APOE, C1QA, C1QB, and C1QC.

Within the stromal compartment, we investigated how ER-driven tumor states might shape surrounding cell populations. The ER⁺ signature showed a strong positive correlation with CAF5 (FAP⁺) fibroblasts (Spearman r = 0.77, P = 2 × 10⁻¹³; Figure 4D) a secretory fibroblast population previously associated with an immune-excluded, “cold” TME^74^. Consistently, CAF5 fibroblasts were significantly more abundant in tumors with low immune infiltration (P = 2.1 × 10⁻¹¹, two-sided Wilcoxon rank-sum test; Figure 4E). Investigating stromal–myeloid coordination we found that CAF5 abundance positively correlated with MC2–C1QA⁺ macrophages (Pearson r = 0.48, P = 5.6 × 10⁻⁴; Figure 3F), linking a stromal population enriched in ER⁺/low-immune tumors to a macrophage subset associated with immunosuppressive signaling.

To further probe mechanisms underlying this coordinated phenotype, we performed ligand– receptor interaction analysis^75^ between macrophages and stromal cells (Methods). Several high-confidence interactions were identified (Figure 4G, Supplementary Table 13), including VIM–CD44 (implicated in stromal remodeling) and TIMP1–CD63 and LGALS3–MCAM (associated with immunoregulatory niches), reinforcing the concept of a stromal–myeloid axis shaping the immune landscape. Together, these findings suggest a functional network in LA breast cancer that links ER⁺ tumor states with immune infiltration, including shifts in myeloid cell populations and the presence of secretory FAP⁺ fibroblasts, ultimately contributing to reduced overall survival and therapeutic resistance.

### EMT-Associated Tumor Programs in TNBC Reveal TROP2 as a Potential Therapeutic Target and Predict Poor Survival In Metastatic lesions

Transcriptional analysis of tumor cells revealed 30 distinct tumor programs (Supplementary Table 8). Investigating their activity level across disease stages (Methods), we found that both TP4 and TP21 were both enriched in TNBC. Both programs are enriched with epithelial-to-mesenchymal transition (EMT)-associated genes, including: SOX9, ATF3, EGR3, multiple heat shock proteins (HSPA1A, HSPA1B, HSPA6, HSP90AA1) and Ubiquitin C ^76–79^. Of note, these programs were active in 65% of TNBC samples, comprising between 16-99% of the total tumor cells in each sample (Figure 5A,B).

**Figure 5:**
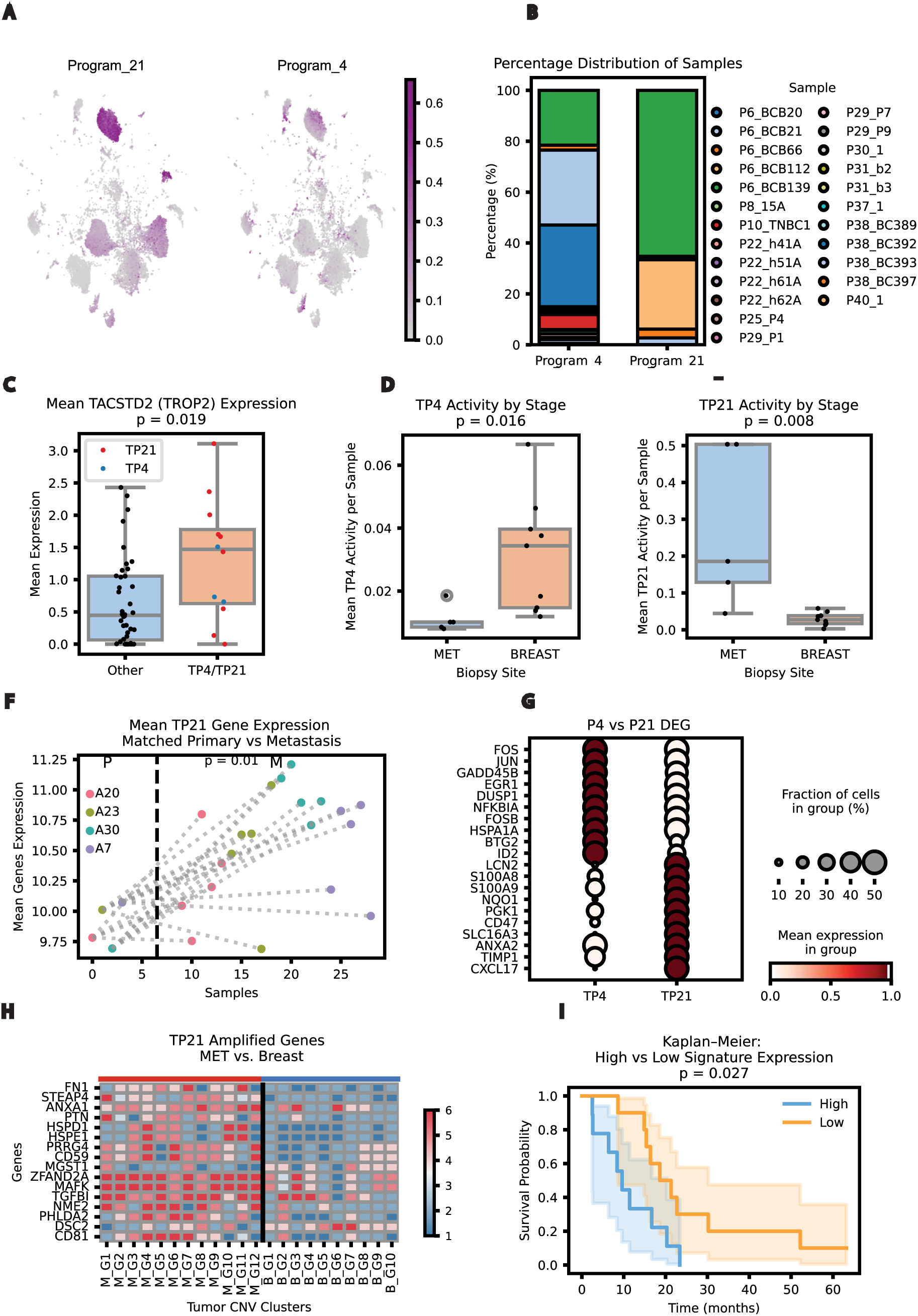
EMT associated tumor programs define transcriptional states, genomic alterations, and clinical outcomes in TNBC. **(A)** UMAP of TNBC tumor cells colored by usage of tumor program 21 (left) and tumor program 4 (right). **(B)** Sample-level distribution of tumor programs 4 and 21. **(C)** TACSTD2 (TROP2) mean expression in tumor cells from program 21 (red), program 4 (blue) vs. all other tumor cells. **(D-E)** Mean activity of (D) TP4 and (E) TP21 across breast cancer biopsy sites. **(F)** Mean expression of TP21 signature genes in primary tumors (left) versus matched metastatic lesions (right). **(G)** Dot plot showing DEGs for tumor program 4 versus tumor program 21.**(H)** Amplified genes in tumor program 21, comparing metastatic (MG1-12) versus primary (BG1-10) CNV clusters. **(I)** Kaplan– Meier survival probability curves comparing TP21-high (>median) versus TP21-low samples (<median).

To further investigate the therapeutic relevance of these transcriptional states, we performed an unsupervised comparison of tumor cells classified under TP4 or TP21 against all other tumor cells. This analysis revealed a significant upregulation of TACSTD2 (encoding TROP2), the therapeutic target of Sacituzumab-govitecan, in cells associated with both programs (p = 0.019, two-sided Wilcoxon rank-sum test; Figure 5C). This finding suggests that tumors exhibiting EMT-like features may be susceptible to TROP2-targeted therapies, currently administrated as a standard line only to metastatic patients.

Examining the difference between these two programs we found that TP4 was significantly higher in primary breast biopsies (P = 0.016, two-sided Wilcoxon rank-sum test, Figure 5D), while TP21 showed significantly elevated activity levels in metastatic lesions (P = 0.008, two-sided Wilcoxon rank-sum test, Figure 5E). To validate the association of TP21 with metastatic progression, we analyzed bulk RNA-seq data from matched primary and metastatic breast cancer lesions profiled by Siegel et al.^80^. We calculated the mean expression of the top differentially expressed genes defining TP21 across each matched primary-metastasis pair (Methods). This gene signature was consistently and significantly elevated in metastases compared to their corresponding primary tumors (P = 0.01, two-sided Wilcoxon rank-sum test; Figure 5F). To explore the transcriptional adaptation of TP21 in the metastatic setting, we compared TP21 with TP4 and identified a significant enrichment of hypoxia-responsive genes in TP21, suggesting a metastasis-associated, adaptive transcriptional state (Figure 5G; Supplementary Table 14). To assess whether this adaptation was driven by genomic alterations, we analyzed the CNV of genomic regions corresponding to the top 100 associated genes in TP21 and TP4. In TP21, we identified amplifications of 16 genes in metastases compared to primary tumors (Figure 5H; Supplementary Table 15; Methods). Conversely, for TP4, we observed amplifications of 8 associated genes in primary tumors relative to metastases CNV clusters (Supplementary Figure 8 Supplementary Table 16; Methods). Furthermore, an independent validation using the AURORA US Network cohort^81^ revealed that high expression of TP21 genes in metastatic disease was associated with significantly worse overall survival (p = 0.027, log-rank test,Figure 5I, Methods). Collectively, these findings suggest that TP21 represents a metastasis-associated transcriptional program linked to poor clinical outcomes and potentially driven by underlying genomic alterations. This program may reflect the evolutionary trajectory of TNBC during disease progression.

### EMT-Driven Tumor States Correlate with Immune Checkpoint Upregulation in TNBC

Focusing on the immune compartment of TNBC tumors, we found that they exhibited significantly higher levels of exhausted T cells compared to other breast cancer subtypes (P = 0.001, two-sided Wilcoxon rank-sum test, Figure 6A). Regulatory T cells from metastatic sites demonstrated increased expression of CTLA4 (P = 0.02, two-sided Wilcoxon rank-sum test, Figure 6B). Conversely, in local disease setting there was elevated co-expression of the immune checkpoints LAG3 and PD1 (P = 0.04, two-sided Wilcoxon rank-sum test, Figure 6C). To validate these findings, we analyzed bulk RNA-seq data from matched primary and metastatic breast cancer lesions profiled by Siegel et al.⁸⁵ and found that PD1 and LAG3 expression was significantly higher in primary tumors than in their matched metastatic sites (P = 0.007, two-sided Wilcoxon rank-sum test, Figure 6D).

**Figure 6.**
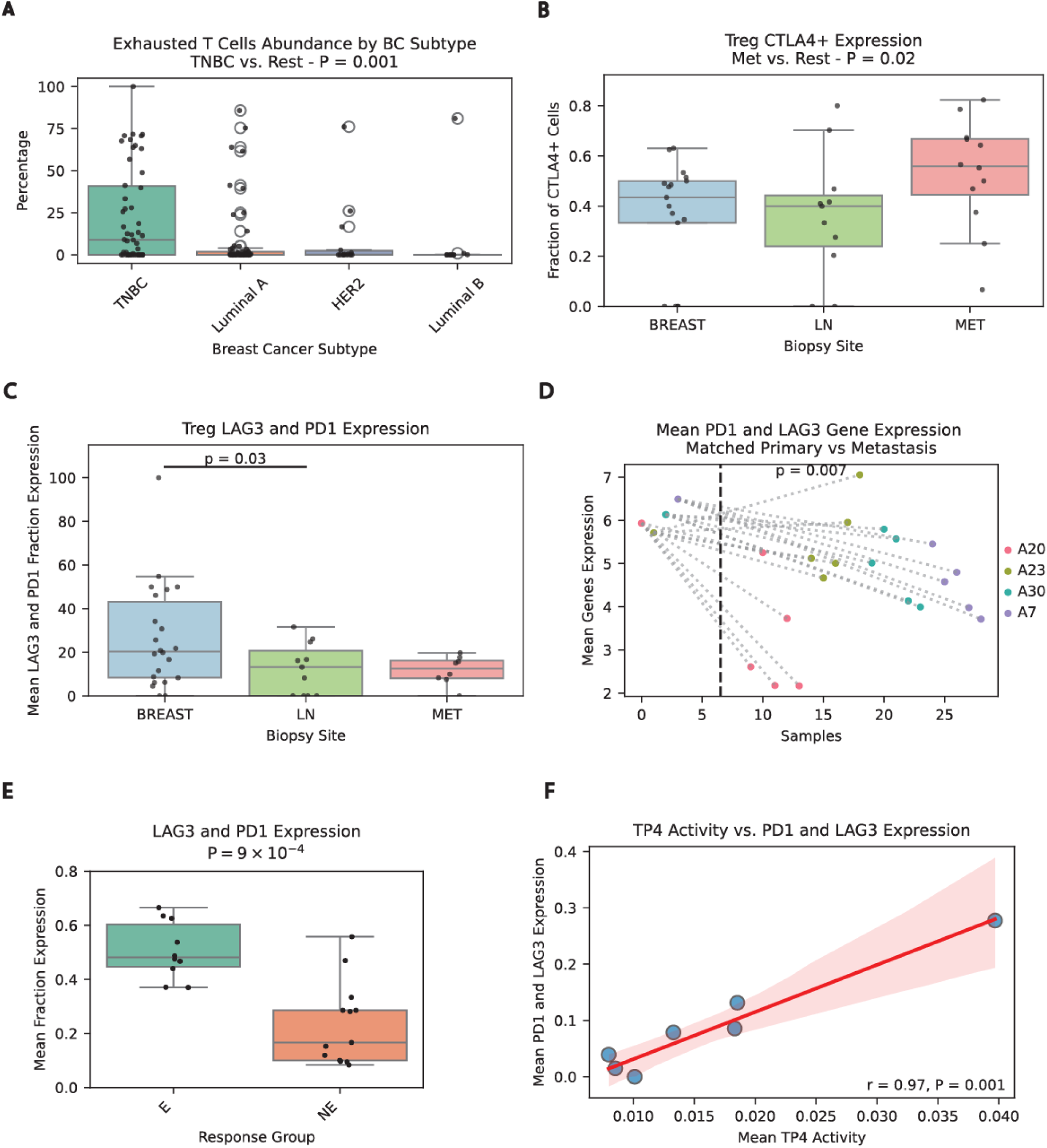
TNBC immune checkpoint landscape correlates with tumor programs. **(A)** Abundance of Exhausted-HS CD8⁺ T cells across breast cancer subtypes. **(B–C)** Fraction of cells expressing (B) CTLA4 and (C) LAG3 and PD1 (average), in regulatory T cells across biopsy sites. **(D)** Mean expression of LAG3 and PD1 (average) in primary tumors (left) versus matched metastatic lesions (right). **(E)** Fraction of cells expressing LAG3 and PD1 in regulatory T cells from expanded (E) versus non-expanded (NE) patients. **(F)** Mean immune cells LAG3 and PD1 correlation with TP4 mean activity level.

Finally, to further contextualize these findings, we examined an independent dataset from Bassez et al., 2021.⁸⁰ In this cohort of BC patients treated with anti-PD1 therapy, patients who experienced T cell expansion exhibited significantly higher expression of LAG3 and PD1 in regulatory T cells compared to those without expansion (P = 9 × 10⁻⁴, two-sided Wilcoxon rank-sum test, Figure 6E). While this observation is consistent with the established role of PD1 and LAG3 as canonical markers of T cell activation and expansion, it highlights the reproducibility of our findings across independent datasets.

As noted above, TP4 exhibited higher activity in primary breast cancers and was positively associated with immune checkpoint expression. Specifically, elevated TP4 scores correlated with increased PD1 and LAG3 expression (r = 0.97, P = 0.001, Methods, Figure 6F). This link between the EMT-like TP4 state and checkpoint upregulation parallels the heightened PD1/LAG3 co-expression observed in local disease, suggesting that TP4-high primary tumors may create an immune-modulatory microenvironment potentially responsive to PD1/LAG3 targeted therapies. While a similar association has been reported in gliomas and other cancer types, to our knowledge, this is the first time it has been demonstrated in breast cancer^82,83^.

Collectively, these findings identify two EMT-associated programs in TNBC with distinct roles: TP4, enriched in primary tumors, is linked to PD-1/LAG3 upregulation and an immune-modulatory microenvironment, while TP21, enriched in metastases, is associated with hypoxia, genomic amplifications, and poor survival. Together, they highlight the coupling of tumor transcriptional states with immune landscape and disease progression.

## Discussion

In this meta-analysis of 31 scRNA-seq studies encompassing over 600,000 cells from 376 samples, we present a comprehensive cellular atlas of the breast tumors microenvironment, integrating immune, stromal, and tumor-intrinsic programs across subtypes and disease stages. Our findings refine the understanding of breast cancer heterogeneity by identifying subtype-specific TME features and transcriptional states that align with the “hot” versus “cold” immune classification— an emerging framework for predicting therapeutic response.

Consistent with prior studies, we confirmed that TNBC exhibits an inflamed TME with high immune infiltration compared to the other BC subtypes. LA tumors exhibited a variably inflamed TME, with primary lesions predominantly displaying a “cold” phenotype—characterized by low immune infiltration, immunosuppressive macrophages, tumor-suppressing chemokine profiles, and secretory CAFs. In metastatic LA tumors, however, we observed greater immune heterogeneity with a subset of patients exhibiting higher immune infiltration and higher abundance of pro-inflammatory macrophages. Our analyses suggest that this immune exclusion/permissive environment is maintained through coordinated tumor-stromal–myeloid interactions.

This immune and stromal heterogeneity is paralleled by transcriptional diversity within tumor cells themselves. Indeed, advances in single-cell technologies have challenged the traditional clinical classification of breast cancer into four molecular subtypes, LA, LB, HER2-enriched, and TNBC, by revealing substantial intragroup heterogeneity not captured by bulk profiling^84^. Considering LA tumors, we here identified two transcriptionally distinct tumor cell populations defined by ER expression status. ER-positive tumor cells demonstrated high expression of proliferation-associated signaling pathways, including KRAS–BRAF–MEK–ERK-MTOR and members of the EGFR family, several of which were found to be associated with chromosomal amplification of the corresponding regions. Although these pathways have previously been associated with ET resistance ^85–87^, our analysis revealed that their activation correlated with improved overall survival and a favorable response to ET, perhaps indicating an additional influence of ET on these pathways as treatment progresses.

Interestingly, HER2 (ERBB2) expression was elevated in ER-positive LA tumors to levels comparable to clinically defined HER2-positive cases. This observation aligns with emerging evidence that HER2-low tumors, despite lacking HER2 DNA amplification, benefit from anti-HER2 therapies, particularly antibody–drug conjugates (ADCs)^64,65^. HER2-low tumors may harbor a higher frequency of HER2 mutations, and HER2 in this context may function either as an oncogenic driver or as a delivery target for cytotoxic payloads via ADCs^88^. Importantly, the ER-positive transcriptional signature was also strongly associated with features of immune exclusion, including low immune infiltration and a cold TME depicted by chemokine, macrophage, and stromal cells phenotypes. This immunologically inert phenotype may contribute to limited responsiveness to immunotherapy in ER-positive tumors^89^.

In contrast, ER-negative LA tumor cells exhibited a distinct metabolic and inflammatory transcriptional program, marked by MYC activation, and enhanced anaerobic glycolysis. This aggressive phenotype was associated with poorer survival outcomes and resistance to ET. However, these tumors also showed increased immune infiltration, suggesting a more immunogenic state that may be responsive to ICB, particularly when combined with conventional chemotherapy or biological agents. Notably, the MYC-driven transcriptional program observed in ER-negative LA tumors highlights a potential therapeutic vulnerability. MYC inhibitors, currently under investigation in early-phase clinical trials, have demonstrated promising preclinical activity, especially in TNBC mouse models^90,91^. These findings raise the possibility that MYC-targeted therapies may offer a novel treatment approach for this high-risk subset of ER-negative LA tumors, for whom standard ET might be less effective.

In TNBC, tumor-intrinsic transcriptional programs underscore the complexity of the TME. TP4, an EMT-associated program enriched in primary tumors, was linked to PD1/LAG3 upregulation and an immune-modulatory microenvironment, whereas TP21, enriched for hypoxia-inducible genes, was more prominent in metastatic lesions and associated with poor overall survival. Despite their aggressive features, both programs displayed elevated TACSTD2 (TROP2) expression, highlighting their potential as biomarkers for TROP2-targeted therapies such as sacituzumab govitecan.

In parallel, we also mapped immune checkpoint differences across disease stages. PD-1 and LAG3 were predominantly expressed in primary tumors, where they have been associated with response to ICB, whereas CTLA-4 expression was elevated in metastatic sites. These findings provide a stage-specific view of immune evasion and may inform the rational design of combination immunotherapy strategies tailored to disease progression.

Our findings offer a comprehensive view of the breast cancer microenvironment; however, several limitations highlight areas for future refinement. First, retrospective integration of datasets may introduce batch effects and potential biases due to differences in sample processing, sequencing depth, and patient selection. While we applied a rigorous preprocessing pipelines and validated findings across multiple cohorts, some residual technical variability may persist. Second, subtype classification relied on metadata annotations and IHC/FISH-derived labels. This classification may vary from tertiary pathology institutes to others. Third, biopsy site variability may confound TME comparisons, particularly in metastatic lesions, where anatomical context influences immune composition. Finally, our analysis was correlative in nature. While we identified associations between gene programs, cell types, and clinical outcomes, functional validation in experimental models is necessary to establish causality, particularly for ligand– receptor pairs mediating immune suppression and for therapeutic predictions based on TROP2 or HER2 expression.

Altogether, our study presents a comprehensive single-cell framework that captures the biological, immune, and stromal heterogeneity of breast cancer across subtypes and disease stages. Through integrative analysis, we identify clinically relevant transcriptional states and genetic signatures, validated in bulk RNA-seq data, that may be applicable in scalable clinical settings. These findings underscore the importance of refined molecular stratification, including the accurate identification of HER2-low and ER-negative subpopulations, to guide treatment decisions and optimize outcomes. Moreover, our analysis of TNBC reveals EMT-driven, metastasis-enriched programs marked by TROP2 expression, supporting the early incorporation of TROP2-targeted therapies in high-risk patients. As single-cell technologies advance, such atlases will be instrumental in translating cellular heterogeneity into therapeutic precision and expanding access to targeted interventions including ICB, ADCs, and MYC inhibitors in the appropriate clinical context.

## Methods

### Preprocessing of Single-cell RNA-seq Datasets

We integrated 31 publicly available scRNA-seq datasets (Supplementary Table 1). All datasets were droplet-based and contained unique molecular identifier (UMI) counts, except for GSE118390, GSE190772 and GSE169246 which provided log₂-transformed normalized expression values. For each dataset, we applied a rigorous preprocessing pipeline as follows: low-quality cells expressing fewer than 300 genes and genes expressed in fewer than three cells were removed. Cells in the top 10% of mitochondrial gene expression within each dataset were excluded, and potential doublets were identified and filtered using the “Scrublet”^92^ algorithm with a ‘doublet rate’ of 0.05. In addition, we eliminated additional doublets by excluding cells that aberrantly co-expressed markers from multiple immune lineages (e.g., CD3E and MS4A1), or tumor cells that expressed canonical immune markers (e.g., CD3E, CD14, CD163, CD19, CD79A, CD8A, CD8B). This quality control steps resulted with 985,551 cells and 10,273 common sequenced genes. All samples were normalized using scanpy ‘scanpy.pp.normalize_total’ function set at ‘target_sum’ of ‘1*e4’, followed by log-transformation: log₂(normalized counts + 1). Finally, when a data set consisted of more than one sample we applied Batch Balanced K-Nearest Neighbors (BBKNN)^93^ with a ‘batch_key’ of ‘sample’ in order to remove batch effects between samples.

### Copy Number Variation Analysis with inferCNV

To distinguish malignant cells from normal epithelial and stromal populations, we performed copy number variation (CNV) analysis on each dataset (excluding immune-only datasets) using the “inferCNV**”** package in R^50^. Immune cells expressing lineage-specific markers (PTPRC, CD3E, CD8A, CD8B, MS4A1, CD14, CD163) were designated as reference cells. Cells without detectable CNVs within the observation group were classified as stromal or normal epithelial. The latter groups were further distinguished by Leiden clustering and their DEG as further elaborated in the-“Dimensionality Reduction and Clustering” section.

### Dataset Integration

Following CNV annotation, cells were grouped into three compartments, tumor, immune, and stroma/normal epithelial, and integrated across all datasets to create three separate merged AnnData objects. In addition, we generated a fourth integrated dataset that included all cells across compartments for global analysis. The integrated dataset that contained all cells as well as each of the three integrated compartments (tumor, immune, and stroma) underwent dimensionality reduction and clustering using the ‘Scanpy’ pipeline^51^: Highly variable genes were identified using ‘scanpy.pp.highly_variable_genes’ with a ‘batch_key’ of ‘sample’, followed by 50 component principal component analysis (PCA). We then applied Batch Balanced K-Nearest Neighbors (BBKNN)^59^ with a ‘batch_key’ of ‘data-set’ in order to remove batch effects between the cohort’s different data sets.

### Clustering and differential expression analysis

Scanpy’s UMAP ‘scanpy.tl.umap’ function was applied for visualization, and Leiden clustering function ‘scanpy.tl.leiden’ was used for community detection. To ensure robustness and avoid dataset-specific artifacts, we required that each cluster be supported by cells from at least two independent datasets. DEGs were identified using Scanpy’s ‘rank_genes_groups’ function with the Wilcoxon rank-sum test. Genes were considered significantly differentially expressed if they met the following thresholds: adjusted p-value < 0.05 and log₂ fold change > 0.5. To validate the results at the sample level, we performed a Wilcoxon rank-sum test on the fraction of cells expressing each gene, defined as those with log₂(normalized counts + 1) > 0.5.

### Ligand–Receptor Interaction Analysis and functional annotation

To identify cell–cell communication signals within the TME, we performed ligand–receptor interaction analysis using the “LIANA” (LIgand-receptor ANalysis frAmework) algorithm^75^. LIANA integrates results across multiple well-established ligand–receptor inference methods—including CellPhoneDB, NATMI, Connectome, and others—and applies a consensus scoring approach. For each dataset (TNBC and Luminal A), the LIANA algorithm was executed three times independently, examining communication in the following directional pairs: immune to stroma, stroma to tumor and tumor to immune. Only ligand–receptor pairs meeting the significance threshold (Cellphone P-value ≤ 0.05) were included. The interactions were visualized as gene-centric chord diagrams, with nodes representing genes and colored by cellular origin (immune = blue, tumor = red, stroma = green). Edges between ligands and receptors were color-coded to reflect their annotated function: red for pro-tumor interactions, blue for anti-tumor, and gray for dual or unknown roles. To provide these annotations we manually curated each significant interaction based on literature evidence. Interactions were classified as tumor-promoters, tumor-suppressors, dual/context-dependent, or unknown according to their known roles in the TME (Supplementary Table 3,4). This classification was guided by published functional studies describing the impact of these signaling axes on tumor progression, immune modulation, angiogenesis, and stromal remodeling.

### ER positive vs. negative signatures

To calculate ER^+^ and ER^−^ gene signatures, we first ranked the DEGs)within each group based on their adjusted p-values (adjusted p < 0.05), from most to least significant. To prioritize genes with potential clinical relevance, we performed survival analysis using the TCGA breast cancer cohort, using normalized bulk RNA sequencing data. Following, the top 150 genes were used to define the ER^+^ signature and the top 50 genes were used to define the ER^−^ signature. Importantly, the results were found to be robust to different selection of gene set size (Supplementary Table 17-18).

### Survival analysis

Kaplan–Meier survival analysis was conducted using the R packages “survival” and “survminer” to assess differences in survival outcomes between patients with high versus low scores for each gene signature. For each patient, survival time and event status (alive\censored or deceased) were derived from the clinical metadata. Survival curves were generated using the survfit function, and statistical significance between groups was evaluated with the log-rank test via the survdiff function (p < 0.05 considered significant).

### CNV Clusters

For each sample, InferCNV clusters cells based on their inferred CNV profiles. Within each cluster, CNV scores are computed across genomic regions, defined by chromosome and gene position. Each gene is assigned a CNV state based on the average expression signal relative to reference cells. According to the InferCNV framework, genes are categorized using a 1–6 numeric scale, where values reflect inferred copy number status (e.g., 1,2 = deletion, 3 = neutral, 5,6 = amplification), as detailed in the InferCNV documentation^50^.

### ER-Based CNV Clusters

Each CNV-defined cluster was evaluated for ER expression by calculating the proportion of ER positive cells within the cluster. Clusters in which more than 50% of cells expressed ER were classified as ER positive CNV clusters, while those with 50% or fewer ER-positive cells were classified as ER negative CNV clusters.

### Identification of Differentially Amplified/Deleted CNV Cluster Genes

To identify genes significantly amplified or deleted between groups of CNV clusters, we performed a Fisher’s exact test for each gene. Specifically, we compared the number of CNV clusters showing amplification or deletion of a given gene in one group vs the other group: e.g., ER-positive vs. ER-negative clusters, or metastatic vs. primary clusters. Genes with a statistically significant difference in amplification or deletion frequency between the groups, based on FDR-adjusted P-values, were considered differentially altered.

### Tumor Program Discovery Using cNMF

To identify recurrent transcriptional states within tumor cells, we applied consensus non-negative matrix factorization (cNMF)^94^ to the integrated tumor cell dataset. This unsupervised algorithm decomposes gene expression data into a set of latent programs (metagenes) and their respective usage (i.e., activity) across cells, allowing the discovery of biologically meaningful gene expression modules. The number of programs (K) was empirically determined by evaluating a range of values for stability and biological interpretability. We selected K = 30 as it balanced two criteria: (1) Reconstruction error: K = 30 minimized error without overfitting, indicating a sufficient number of programs to capture key transcriptional patterns; (2) Biological resolution: This value preserved distinct, non-redundant gene programs (e.g., EMT, hypoxia, cell cycle), while avoiding fragmentation into noisy or overlapping components seen at higher K values. The cNMF output included: A usage matrix, denoting the degree to which each program contributes to the transcriptome of each cell, and a gene loading matrix, listing the top genes defining each program. These programs were further characterized by pathway enrichment and biological themes, and subsequently used in comparative analyses across breast cancer subtypes, response groups, and metastatic progression.

### Validation analysis

5 independent cohorts were used for validation of our transcriptomic results:

1. TCGA Cohort Survival Analysis^72^: The survival plots for both ER positive and ER negative signatures was described above. To further validate the results, an empirical p-value was calculated by repeating the survival analysis 100 times using the mean expression of 150, 50 randomly selected genes. For each iteration, the number of times (n) the resulting p value was more significant than the original observed value was recorded. For both ER positive and ER negative signatures, the original p-value remained more significant than (n + 1) / 101 supporting the robustness and statistical significance of the findings. To determine whether these signatures provide prognostic value independent of established clinical variables, we performed multivariable Cox proportional hazards regression analyses. For both ER-negative and ER-positive signatures, the clinical variables considered in the model were: age, stage (TNM), menopause status, ethnicity, and surgical margin status (Supplementary Tables 17,18)
2. Xia, et al. 2022^73^-bulk raw RNA sequenced counts were downloaded from EMBL-EBI ArrayExpress database under the accession number E-MTAB-9917. The data was normalized followed by log₂(normalized counts + 1)conversion. For each sample the mean of the ER positive and ER negative signatures was calculated and compared between treatment groups. Importantly, the results were found to be robust to different selection of gene set size (Supplementary Tables 19,20).
3. Bassez et al. 2021^95^-the single-cell RNA raw counts were downloaded from <u>EGAS00001004809</u>. The data was then processed using our pipeline for single cell data detailed above. Metadata for patient’s response and cell annotations were merged. For each sample the mean fraction expression of PD1 and LAG3 was calculated.
4. Siegel et al. 2018^80^-Normalized and log-transformed bulk RNA sequencing data were obtained from GSE110590. For each sample, the mean expression of the top 30 genes from TP21 was calculated, excluding samples in which the primary tumor had been treated with chemotherapy and radiotherapy. A Wilcoxon rank-sum test was used to assess differences in TP21 expression between metastatic and matched primary tumor samples. The results were found to be robust to various gene set size (Supplementary Table 21)
5. Aurora US cohort^81^ Survival Analysis - bulk raw RNA sequenced counts were downloaded from Aurora website including patient’s metadata. The data was normalized and log transformed as was described in Xia et al. publication^73^. TNBC patients were selected from the dataset. Kaplan–Meier survival analysis using the lifelines Python package^96^ was applied by stratifying patients into high/low groups using the top 30 genes of TP21. Importantly, this result was found to be robust to different gene set sizes (Supplementary Table 22).

### TP4 correlations

The mean TP4 activity of each sample was correlated with the mean PD-1 and LAG3 expression in immune cells, excluding one outlier sample with TP4 activity levels fourfold higher than the cohort mean.

## Author contributions

L.Z.S and K.Y. conceived the idea. L.Z.S. and K.Y. designed the study. L.Z.S performed the analysis. A.P. performed the cNMF analysis and created the manuscript-related website. L.Z.S and K.Y. wrote the manuscript.

## Materials & Correspondence

This study did not generate new unique reagents.

